# Hepatoprotective effects of *Erythrina abyssinica* Lam Ex Dc against Non Alcoholic Fatty Liver Disease in Sprague Dawley Rats

**DOI:** 10.1101/577007

**Authors:** Felix Kanyi Macharia, Peter Waweru Mwangi, Abiy Yenesew, Frederick Bukachi, Nelly Murugi Nyaga, David Kayaja Wafula

## Abstract

**Background:** Non-alcoholic fatty liver disease (NFLD) is the hepatic manifestation of the metabolic syndrome recognized as the most prevalent chronic liver disease across all age groups. NFLD is strongly associated with obesity, insulin resistance, hypertension and dyslipidemia. Extensive research efforts are geared, through pharmacological approach, towards preventing or reversing this. *Erythrina abyssinica* Lam ex DC is an indigenous tree used widely used in traditional medicine, including for the treatment of liver related diseases, and has been shown to possess hypoglycemic, anti-oxidant, antimicrobial and anti-plasmodia effects. The present study is aimed at establishing the effects of *E. abyssinica* on the development of Non-alcoholic fatty liver disease induced by a high-fat and high-sugar diet in rats, *in-vivo* model.

**Methods:** Forty rats (40) were randomly divided into five groups: positive control (pioglitazone), Negative control (high fat/high sugar diet), low test dose (200 mg/kg), high test dose (400 mg/kg) and normal group (standard chow pellets and fresh water).

The inhibitory effect of the stem bark extract of *E. abyssinica* on the development of NAFLD was evaluated by chronic administration the herb extracts to rats on a high-fat/high-sugar diet. Biochemical indices of hepatic function including serum lipid profile, serum aspartate transaminase and alanine transaminase levels were then determined. Histological analysis of liver samples was carried out to quantify the degree of steato-hepatitis. Liver weights were taken and used to determine the hepatic index. The data was analyzed using one-way ANOVA, and Tukey’s post-hoc tests were done in cases of significance. Histology data was analyzed using Kruskal-Wallis and Dunn’s post-hoc test was done in cases of significance. Significance was set at p<0.05.

**Results:** The freeze dried extract of *E. abyssinica* had significant effects on **fasting blood glucose** [5.43 ± 0.17 (HF/HSD) vs 3.8 ± 0.15 (E 400 mg/kg) vs 4.54 ± 0.09 (E 200 mg/kg) vs. 4.16 ± 0.13 (PIOG) vs. 2.91 ± 0.16 (normal control): P value < 0.0001], and **insulin sensitivity** [329.4 ± 13.48 mmol/L · min (HF/HSD) vs. 189.8 ± 12.11 mmol/L · min (E 400 mg/kg) vs. 233.8 ± 6.55 mmol/L· min (E 200 mg/kg) vs. 211.1 ± 7.35 mmol/L · min (PIOG) vs. 142.9 ± 11.94 mmol/L · min: P value < 0.0001],

The extract had significant effects on hepatic indices including, **hepatic triglycerides** (P value < 0.0001), **liver weights** (P value < 0.0001), **liver weight-body weight ratio** (P value < 0.0001), **serum ALT levels** (P value < 0.0001), **serum AST** (P value < 0.0017), **serum total cholesterol** (P value < 0.0001), **serum triglycerides** (P value < 0.0001), and **serum LDL-cholesterol** (P value < 0.0001). The extracts however showed no significant effects on **HDL-cholesterol** (P value = 0.4759).

Histological analysis showed that the extract appears to possess protective effects against steatosis, inflammation and hepatic ballooning, with the high dose (400mg/kg) being more hepato-protective.

**Conclusion:** The freeze dried stem bark extract of *Erythina abyssinica* possesses significant inhibitory effects against the development of NAFLD in Sprague Dawley rats.

## 1.0 Introduction

Non-alcoholic fatty liver disease (NAFLD) is a condition characterized by excessive accumulation of hepatic fat (primarily triacylglycerols) that occurs in the absence of secondary causes such as alcohol consumption, use of steatogenic medication and steatogenic hereditary disorders[1–4]. This condition which has a prevalence of between 6% and 35% globally may often progress to hepatic failure or hepatocellular carcinoma and is the leading cause of chronic liver disease worldwide [2,3].

Although initially recognized as being primarily as a hepatic disorder, NAFLD is now known to affect multiple organs in the body such as the heart, kidneys and endocrine organs, and is a major risk factor in chronic diseases affecting these organs [5–8]. More pertinently, NAFLD is now recognized as the hepatic manifestation of metabolic syndrome with the majority of NALFD patients having some, if not all, features of this syndrome [2,9,10]. Indeed, longitudinal studies suggest that NAFLD precedes development of both type 2 diabetes mellitus (T2DM) and metabolic syndrome [6,8,11]. Unfortunately, no pharmacological treatment has been shown to reverse the natural history of NAFLD with current treatment regimens focusing mainly on exercise and dietary modifications [12,13].

This study investigated the effects of the chronic administration freeze dried extracts of *Erythrina abyssinica* Lam ex DC (a plant species which has several medicinal uses in African communities and possesses demonstrated hypoglycemic and cytotoxic effects) in preventing the development of NAFLD in Sprague Dawley rats. The NALFD was induced by chronically feeding the rats a high fat/high sugar diet containing monosodium glutamate.

The freeze-dried stem bark extracts of *Erythrina abyssinica* Lam Ex DC significantly decreased biochemical and histological markers associated NAFLD namely: fasting blood glucose, insulin resistance, serum triglycerides, serum total cholesterol, serum ALT, hepatocellular ballooning, micro-vesicular steatosis and lobular inflammation. The freeze-dried extracts of *Erythrina abyssinica* Lam ex. DC therefore significantly inhibited the development of NAFLD in Sprague Dawley rats.

## 2.0 Materials and Methods

The study was conducted in the Department of Medical Physiology, University of Nairobi.

### 2.1 Plant Collection and Extract Preparation

The stem barks of *Erythrina abyssinica* Lam ex DC were harvested from trees growing in Kitui County, Kenya and the identity of the collected plant specimens verified by the staff of the university herbarium located at the School of Biological Sciences, University of Nairobi and a voucher specimen deposited therein (EA2016). The plant parts were air dried for ninety-six hours after which they were milled into a fine powder at the Department of Chemistry, University of Nairobi.

The weight of the resultant powder was then measured using an analytical scale. The powder was then macerated in water in a ratio of 1:10 (weight: volume) and left to stand for one (1) hour. The resulting suspension was then sequentially filtered using cotton wool and Whatmann’s® filter paper to obtain the filtrate which was then placed in a standard laboratory freezer overnight. The filtrate was then lyophilized, and the resulting freeze-dried extract weighed and placed in amber colored sample bottles and stored within a laboratory freezer until when required for administration to the experimental animals.

### 2.2 *In vivo* experiments

Forty (40) freshly-weaned (4 week old) Sprague-Dawley rats (twenty female and twenty male) weighing between thirty to fifty grams were randomized into the normal control (normal diet and distilled water), negative control (high fat/ high sugar diet only), the positive control (high fat/high sugar diet plus pioglitazone), low dose test (high fat/ high sugar diet plus 200mg/kg freeze dried extract of *Erythrina abyssinica* and high dose test (high fat/high sugar diet plus 400mg/kg freeze dried extracts of *Erythrina abyssinica*) groups.

The experimental animals were housed at the animal house located within the Department of Medical Physiology University of Nairobi. The following were the conditions maintained within the animal house: at a room temperature of 23.2°C, relative humidity of 60+/- 10 with twelve-hour light/dark cycles. They were habituated for two weeks prior to the start of the study and were fed standard chow pellets ad libitum during this period.

The high fat/ high sugar diet was prepared by the addition of 20% w/w saturated fat (Cowboy™ manufactured by Bidco™ industries) to standard rat chow which contained protein 29.82%, fat 13.43% carbohydrates 56.74%, fiber 5.3%, vitamins and minerals small quantities-ppm (Unga™ Feeds) to ensure that the pellets delivered approximately 30% of calories as fat. Sucrose on the other hand was added to drinking water to make up a 30% sucrose solution. Monosodium glutamate (MSG) was added to the pellets to make up 0.8% of the pellets to increase the palatability of the pellets [14]. The resulting high-fat/high sugar diet contained 30% fat and 30% sucrose in water.

All the animals received diets ad libitum and the extracts/treatments were administered via oral gavage.

### 2.3 Ethical approval

This study was carried out in strict accordance with the recommendations in the Guide for the Care and Use of Laboratory Animals of the National Institutes of Health. The protocol was approved by the bio-safety, animal use and ethics committee Department of Veterinary physiology, University of Nairobi (REF: FVM BAUEC/2016/111) In addition, the experimental design obeyed the 4Rs (reduction, replacement, refinement and rehabilitation) of ethical animal experimental design [15,16].

### 2.4 Assay of fasting blood glucose

Assay of fasting blood glucose was performed in the manner described by Lee and Goossens [17]. Briefly blood was sampled from the lateral tail vein after a six hour fast. The tail was warmed by dipping in warm water for five (5) minutes to cause vasodilation of the vein and therefore ease visualization and blood sample collection. The blood glucose was determined using a glucometer (Nova Biomedical) and recorded.

### 2.5 Insulin tolerance test

Insulin tolerance tests were performed on Days 21, 42 and 56. The insulin tolerance tests were performed according to the protocol by Bowe [18]. Briefly the experimental animals were fasted for six (6) hours. They were thein weighed after which soluble insulin (Humulin S™ Eli Lilly USA) was administered by an intraperitoneal injection at a dose of 0.75IU/kg. The blood glucose levels were then assayed at regular time intervals after the administration of the insulin as described in 3.2.2. (at time 0 minutes, 30 minutes, 60 minutes, 90 minutes). A 50% Glucose solution was made available to prevent fatalities due to insulin-induced hypoglycemia (i.e. if blood glucose levels ≤1.1 mmol/L). The animals would be withdrawn from the procedure if this happened and the glucose solution administered via oral gavage. The areas under the curve (AUCs) for the respective rats were then calculated and then recorded.

### 2.6 Assay of serum biochemistry

The procedure for determination of the serum biochemical parameters was as follows: the rats were fasted overnight on the last day of the study (day fifty-six) and then sacrificed under euthanasia using pentobarbital. The absence of the Pupillary light reflex was used as a confirmative marker of death. Five (5) milliliters of blood were harvested from each using cardiac puncture. The collected blood was then centrifuged to obtain serum which was then assayed for the following biochemical parameters: fasting serum triglycerides, HDL cholesterol, LDL-cholesterol and total cholesterol, aspartate transaminase (AST) land alanine transaminase. The assays were performed at the Clinical Chemistry laboratory, Kenyatta National Hospital.

### 2.7 Assay of hepatic triglycerides

A midline abdominal incision was performed after the rats were euthanized and the liver tissues excised. The respective livers were then weighed using an analytical balance and the weights recorded. The respective liver tissues were each divided into two unequal portions with the larger portion weight: smaller portion weight ratios being 2:1. The smaller portions were used in the hepatic triglyceride assay while the larger portions were used in the histological studies.

The smaller portions were stored in a laboratory freezer and then later homogenized with phosphate buffer acting as the liquid medium. The resulting liquid homogenates were then used to determine hepatic triglyceride content using the protocol previously described by Butler et.al [19].

### 2.8 Liver histology

The larger portions of the harvested liver tissues were preserved in 10 % neutral formaldehyde. Sections (4 mm^2^ in size) were later fixed in PAF and embedded in paraffin. These sections were further sliced to 5 µm and stained with H/E stain for evaluation of hepatic steatosis.

Liver tissues were scored according to the grading system introduced by [20] for NAFLD: Steatosis: Grade 1 < 33 % steatotic hepatocytes, Grade 2 < 33% to 66% steatotic hepatocytes, grade 3 > 66% steatotic hepatocytes in a X 200 microscopic field of view.

Hepatocellular ballooning: Grade 1-minimal zone 3, Grade 2 – present zone 3, Grade 3-marked predominantly zone 3.

Lobular inflammation: 0-absent, grade 1 <2 foci of inflammation, grade 2: 2-4 foci, grade 3 > 4 foci.

Portal inflammation 0 –absent, grade 1 – mild, grade 2-moderate, grade 3-severe.

Stages of fibrosis: stage 1: zone 3 peri-sinusoidal fibrosis, stage 2-focal or extensive periportal fibrosis, stage 3-bridging fibrosis, stage 4-cirrhosis.

### 2.9 Statistical analysis

The experimental data were expressed as Mean ± S.E.M. and analyzed using One-way ANOVA and Tukey’s post-hoc tests carried out in cases of significance (set at p<0.05) using Graph Pad Prism™ software version 6. The semi-quantitative histological data were analyzed using the Kruskal-Wallis and Dunn’s post-hoc tests were carried out in cases of significance which was also set at p<0.05.

## 3.0 RESULTS

### 3.1 Extract preparation

The stem barks of *Erythrina abyssinica* Lam ex DC were harvested from trees growing in Kitui county, Kenya. The plant parts were air dried for forty-eight hours. The dried parts were then milled at the Department of Chemistry, University of Nairobi. The weight of the resultant powder was measured using an analytical scale. The powder was mixed in a ratio of 1:10 with water in a flask and swirled severally before being left to settle. The mixture was then filtered using cotton wool and Watmann® filter paper to obtain a crude extract solution which was frozen and lyophilized at ICIPE Nairobi, Kenya.

The percentage yield of the extract was 3.4% (885.3g of plant powder yielded 29.88g of extract)

### 3.2 Fasting blood glucose results

There were significant differences in blood glucose levels between the experimental groups on day seven (7) [4.89 ± 0.24 mmol/L (negative control) vs 4.01± 0.21 mmol/L (test high dose) vs 4.14 ± 0.14 mmol/L (test low dose) vs. 4.13 ± 0.19 mmol/L (positive control) vs 3.56 ± 0.30 mmol/L (normal control): P value = 0.0043]. Post-hoc analysis showed that there were significant differences between the negative control and normal control groups (P value = 0.0017).

There were also significant differences between the experimental groups on day fourteen [5.19 ±0.24 mmol/L (negative control) vs. 4.03 ± 0.30 mmol/L (test high dose) vs. 4.7 ± 0.20 mmol/L (test low dose) vs. 4.26 ± 0.20 mmol/L (positive control) vs. 3.83 ± 0.40 mmol/L (normal control): P value = 0.01]. Post-hoc analysis showed that there were significant differences between the negative control and the high dose test groups (P value = 0.0402) as well as between the negative control and the normal control groups (P value = 0.0012).

There was also significant difference between the experimental groups on day twenty-one [5.37 ± 0.27 mmol/L (negative control) vs. 4.38 ± 0.15 mmol/L (test high dose) vs. 4.95 ± 0.24 mmol/L (test low dose) vs. 4.8 ± 0.16 mmol/L (positive control) vs. 3.75 ± 0.21 mmol/L (normal control): P value < 0.0001]. Post-hoc analysis showed significant differences between the negative control and high dose test groups (P value = 0.017). Significant differences were also observed between the normal control and low dose test groups (P value = 0.0025) as well between the normal control and the negative control groups (P value < 0.0001).

There were significant differences between the experimental groups on day twenty-eight [5.3 ± 0.21 mmol/L (negative control) vs. 4.39 ± 0.14 mmol/L (test high dose) vs. 4.48 ± 0.29mmol/L (test low dose) vs. 4.49 ± 0.13mmol/L (positive control) vs. 3.75 ± 0.13mmol/L (normal control): P value < 0.0001]. Post-hoc analysis showed that there were significant differences between the negative control and high dose test groups. (P value = 0.0166). In addition, there were significant differences between the two test groups (i.e. low dose and high test) (P value = 0.0407). As previously, there were significant differences between the negative control and normal control groups high-fat/high sugar diet groups (P value < 0.0001).

There were significant differences between the experimental groups on day thirty-five [4.89 ± 0.24mmol/L (negative control) vs. 4.01 ± 0.21mmol/L (test high dose) vs. 4.13 ± 0.14mmol/L (test low dose) vs. 4.13 ± 0.19mmol/L (Positive control) vs. 3.56 ± 0.31 (normal control): P value < 0.0049]. Post-hoc analysis showed significant differences between the negative control and the high dose test groups (P value = 0.0136) as well as between the negative control and positive control groups (P value = 0.0027).

There were significant differences between the experimental groups on day forty-two [5.26 ± 0.21mmol/L (negative control) vs. 3.63 ± 0.13mmol/L (test high dose) vs. 4.06 ± 0.11mmol/L (test low dose) vs. 4.23 ± 0.17mmol/L (positive control) vs. 2.15 ± 0.08mmol/L (normal control group): P value < 0.0001]. Post-hoc analysis showed significant differences between the negative control and high dose test groups (P < 0.0001) as well as between the negative control and the low dose test groups (P < 0.0001). There were however no significant differences between the low dose test and the high dose test groups (P value = 0.2465) as well as between the high dose and positive control groups (P value = 0.0508).

There were significant differences between the experimental groups on day forty-nine [5.27 ± 0.19mmol/L (negative control) vs. 3.74 ± 0.08mmol/L (test high dose) vs. 4.38 ± 0.10mmol/L (test low dose) vs. 3.73 ± 0.13mmol (positive control) vs. 2.68 ± 0.15mmol/L (normal control): P value < 0.0001]. Post-hoc analysis showed there were significant differences between the negative control and high dose test groups (P < 0.0001) as well as between the negative control and the low dose test groups (P < 0.0001). There were also significant differences between the negative control and normal control groups (P < 0.0001).

There were significant differences between experimental groups on day fifty-six [5.43 ± 0.17mmol/L (negative control) vs. 3.8 ± 0.15mmol/L (test high dose) vs. 4.54 ± 0.09mmol/L (test low dose) vs. 4.16 ± 0.13mmol/L (positive control) vs. 2.91 ± 0.16 (normal control): P < 0.0001]. Post hoc analysis showed there were significant differences between the negative control and the high dose test groups (P < 0.00001) as well between the negative control and the low dose test groups (P < 0.0001). There were significant differences between the two test groups (P value = 0.0070). In addition, there were significant differences between the normal control group and either of the test groups i.e. the low dose test (P < 0.0001) and the high dose test (P = 0.0009) groups. There were however, no significant differences between the positive control and either of the test groups i.e. the high dose test (P = 0.3918) or low dose test (P = 0.3580) groups.

The results of the fasting glucose experiments are graphically represented in figures 1 (A-H).

**Figure 1.**
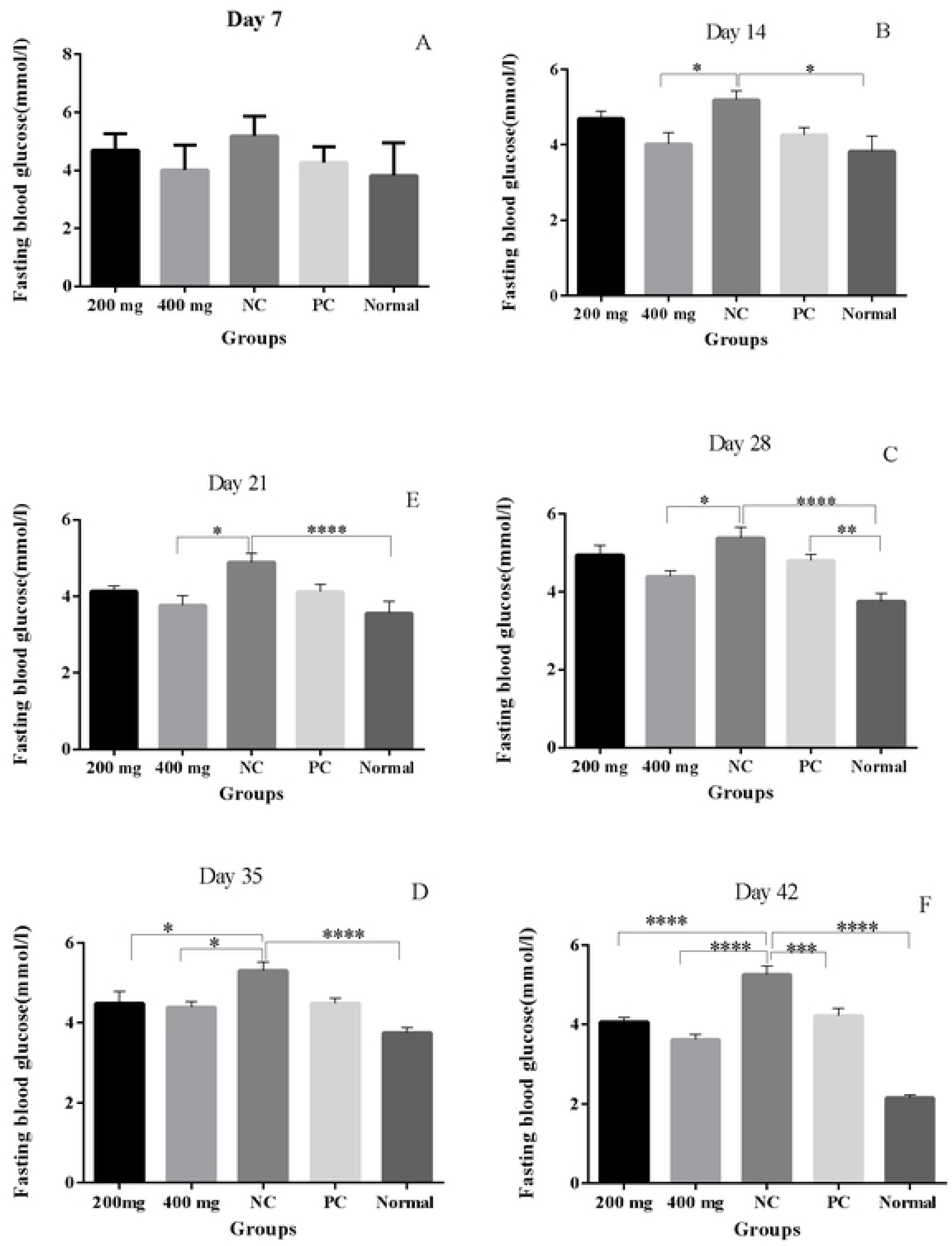
**(A-E)** Fasting blood glucose in mmol/l among the five groups, on day 7(A), day 14(B), day 21 (C), day 28 (D), day 35 (E), day 42 (F), day 49 (G), and day 56(H). * P VALUE < 0.05, **P VALUE <0.01, ***P VALUE <0.001, **** P VALUE<0.0001.

### 3.3 Insulin Tolerance Test results

#### (i) Day 21

There were significant differences in the area under the curves (AUCs) of the insulin tolerance tests between the experimental groups on day twenty-one [237.9 ± 21.39 mmol/L· min (negative control) vs. 179.1 ± 16.55 mmol/L · min (test high dose) vs. 199.3 ± 13.12 mmol/L · min (test low dose) vs. 230.6 ± 17.79 mmol/ L · min (positive control) vs. 116.3 ± 12.15 mmol/L · min (normal control): P < 0.0001].

Post-hoc analysis showed significant differences in insulin sensitivity between the negative control and normal control (P value < 0.0001) groups as well as between the normal group and the low dose test groups (P = 0.0092). There were also significant differences between the normal control and the positive control groups (P = 0.0239). A graphical presentation of these experimental data is shown in figure 2.

**Figure 2.**
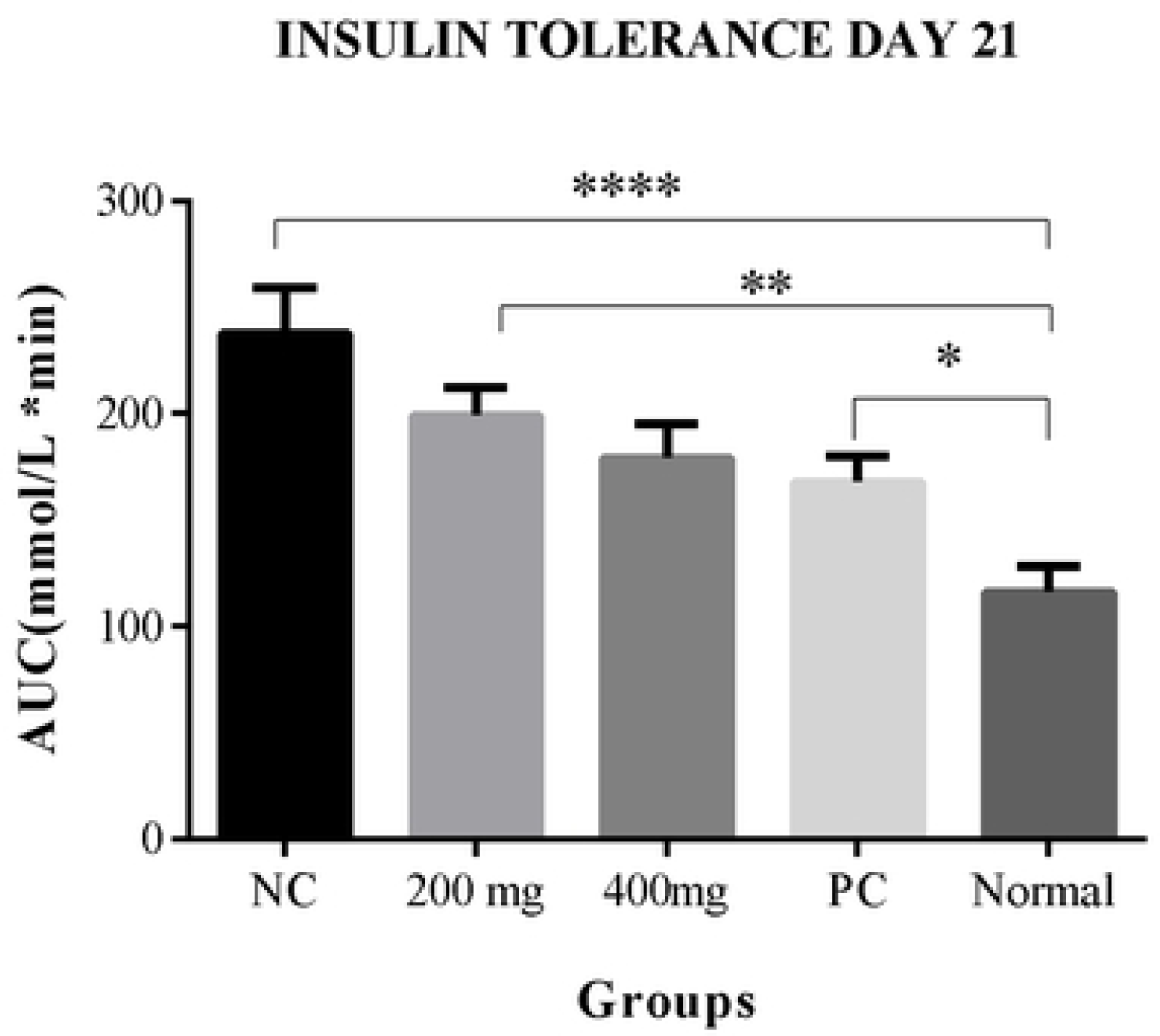
Area under the curve for insulin tolerance tests in mmol/L·min at day 21*P value < 0.05, **P value < 0.01, ****P value < 0.0001

#### (ii) Day 42

There were significant differences in the area under the curves (AUCs) of the insulin tolerance tests between the experimental groups on day forty-two [332.8 ± 37.84 mmol/L · min (negative control) vs. 180 ± 7.48 mmol/L · min (test high dose) vs. 258.4 ± 9.29 mmol/L · min (test low dose) vs. 186.6 ± 7.1 mmol/L · min (positive control) vs. 111.4 ± 4.66 mmol/L · min (normal control): P < 0.0001].

Post hoc analysis showed significant differences in insulin sensitivity between the negative control and the high dose test (P < 0.0001) as well as the low dose test (P value < 0.0471) groups. There were however significant differences between the two test groups (i.e. low dose and high dose) (P = 0.0399). There were significant differences between the normal control and low dose test groups (P < 0.0001) as well between the normal control and the high dose test groups (P = 0.0760). As would be expected, there were significant differences between the negative control and normal control (P < 0.0001).

A graphical representation of the experimental data is shown in figure 3

**Figure 3.**
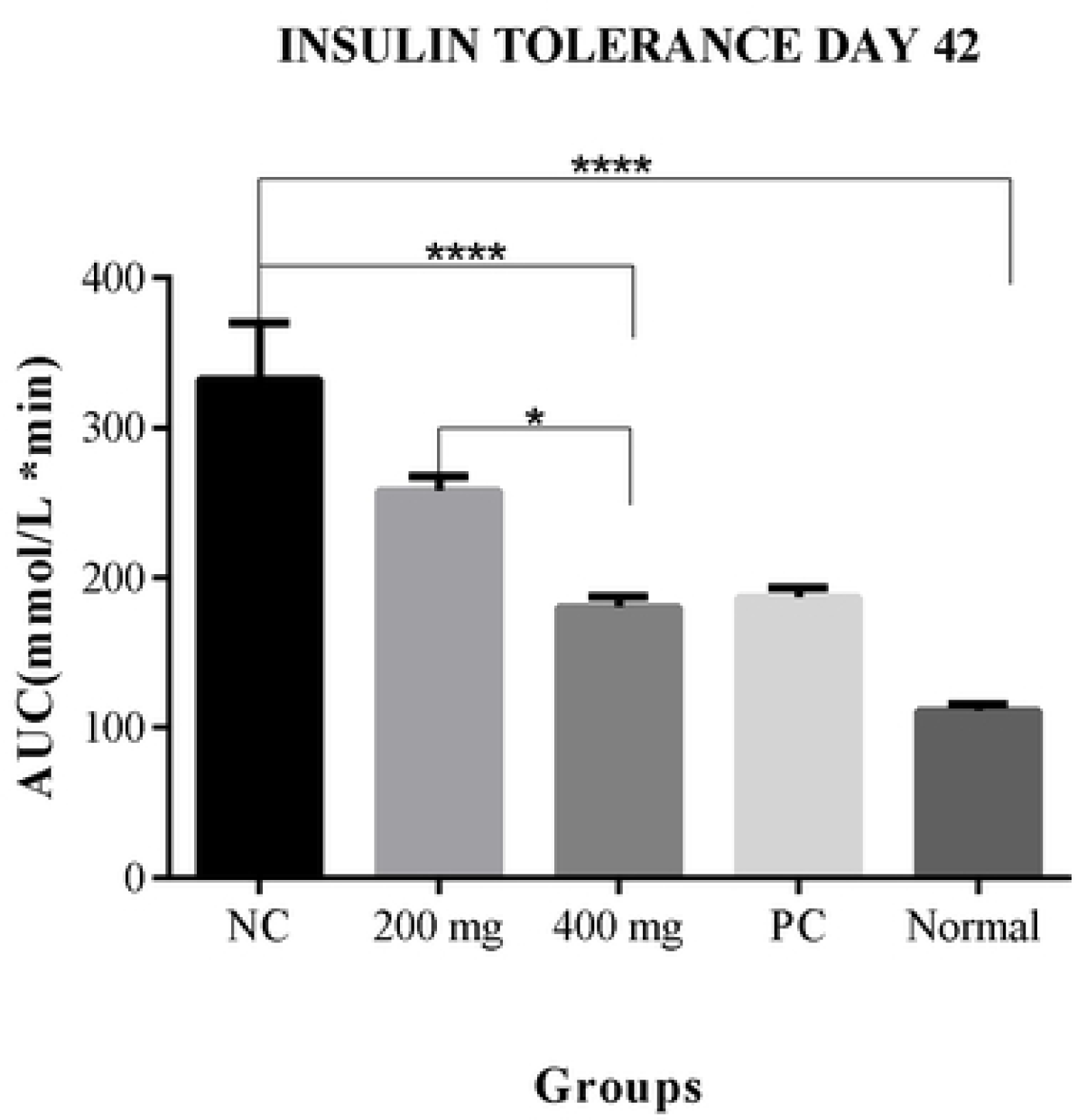
Area under the curve for insulin tolerance tests in mmol/L·min at day 42. *p<0.05, **p<0.01, ****p<0.0001.

There were significant differences in the area under the curves (AUCs) of the insulin tolerance tests between the experimental groups on day fifty-six [329.4 ± 13.48 mmol/L · min (negative control) vs. 189.8 ± 12.11 mmol/L · min (test high dose) vs. 233.8 ± 6.55 mmol/L · min (test low dose) vs. 211.1 ± 7.35 mmol/L · min (positive control) vs. 142.9 ± 11.94 mmol/L · min (normal control): P < 0.0001].

Post-hoc analysis showed significant differences in insulin sensitivity between the negative control and the high dose test (P < 0.0001) as well as the low dose test (P < 0.0001) groups. There were significant differences between the high dose test and low dose test groups (P= 0.0447). There were however also significant differences between the normal control and the high dose test (P value = 0.0286) and the low dose test groups (P < 0.0001).

There were no significant differences between the positive control and the low dose test (P= 0.5635) or the high dose test (P = 0.6230) groups. A graphical representation of the experimental data is shown in figure 4

**Figure 4.**
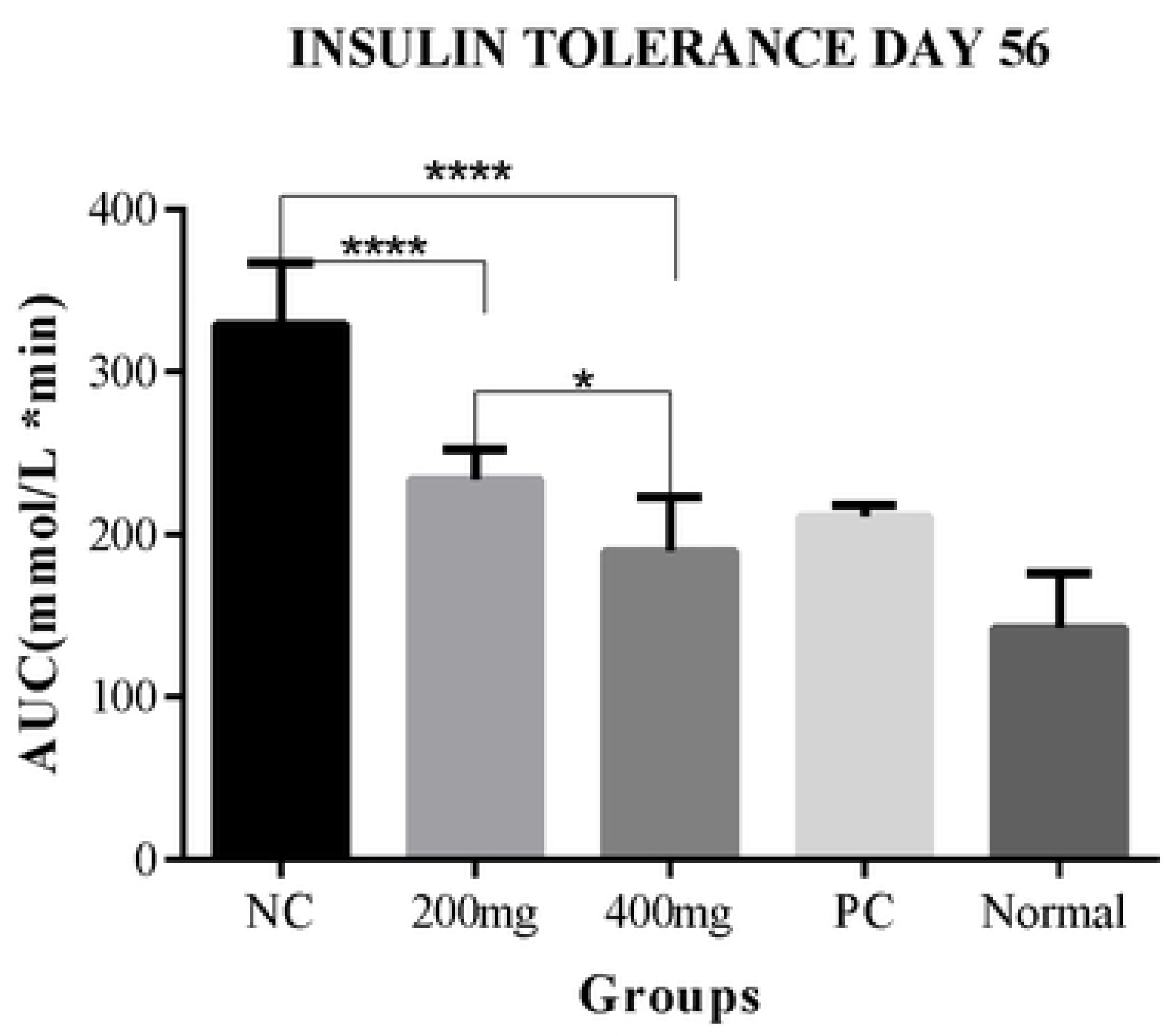
Area under the curve for insulin tolerance tests in mmol/L·min at day 56. *P value < 0.05, **P value < 0.01, ****P value < 0.0001.

### 3.4 Hepatic Triglycerides

There were significant differences in the mean hepatic triglyceride levels between the experimental groups [6.09 ± 0.56 mg/gram of liver tissue (negative control) vs. 2.06 ± 0.21 mg/gram (test high dose) vs. 3.86 ± 0.55 mg/g (test low dose) vs. 3.92 ± 0.44 mg/ gram (positive control) vs. of 2.05 ± 0.28 mg/gram (normal control): P < 0.0001]. Post-hoc analysis showed significant differences in the levels of hepatic triglycerides between the negative control and normal control groups (P value < 0.0001). There were significant differences between the negative control and both the high dose test (P<0.0001) and low dose test groups (P=0.0071). There were significant differences between the low dose and high dose test groups (P = 0.0414).

The experimental data are graphically represented in figure 5.

**Figure 5:**
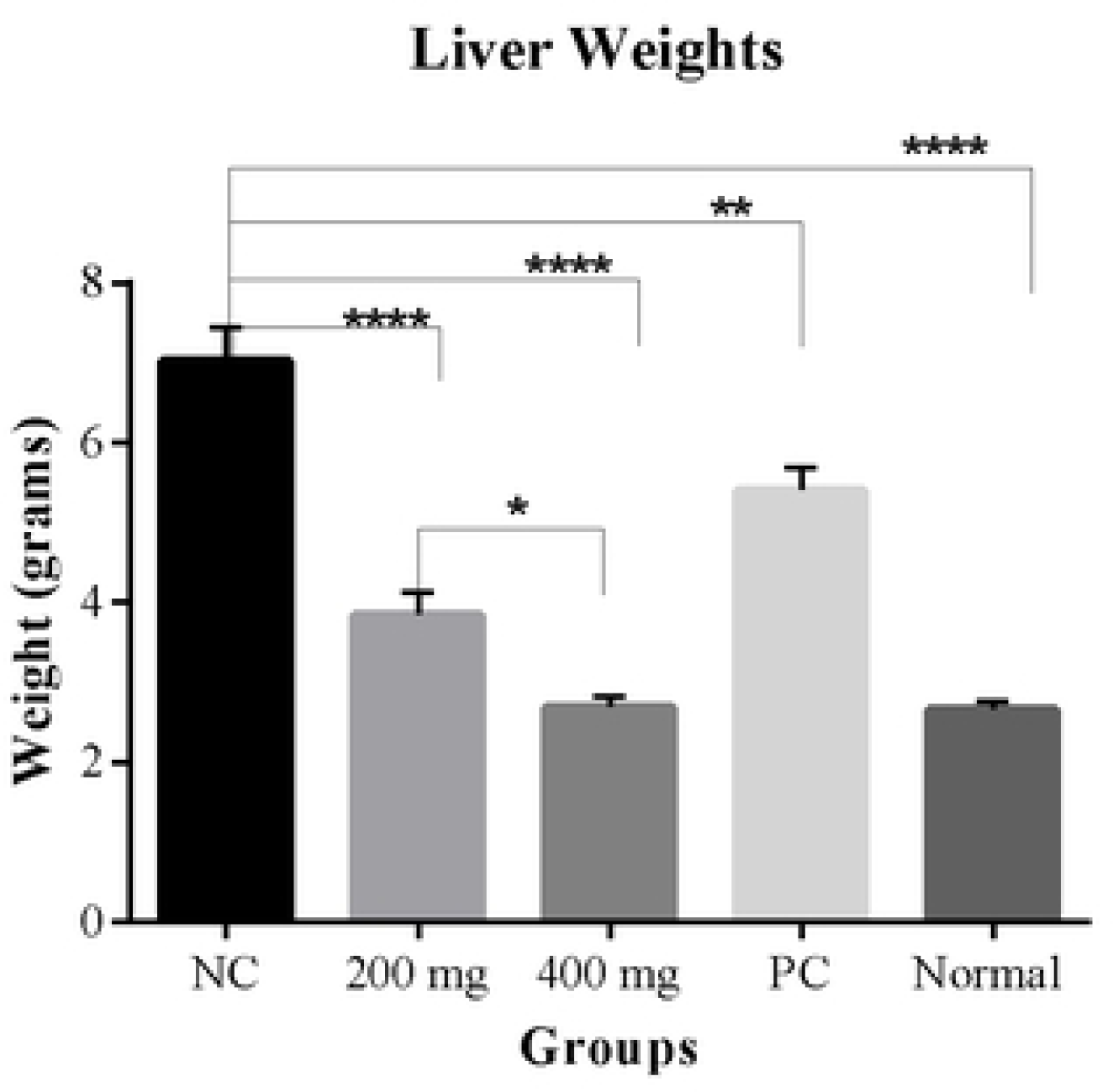
Hepatic triglycerides in mmol/L. NC (Negative control), PC (Positive control) **P value<0.01****P value < 0.0001.

### 3.5 Hepatic weights

There were significant differences in liver weights between the five groups [7.02 ± 0.42 g (negative control) vs. 2.68 ±0.14 g (high dose test) vs. 3.84 ± 0.29 g (low dose test) vs. 5.40 ± 0.29 g (Positive control) vs. 2.65 ± 0.11 g (normal control): P value < 0.0001]. Post-hoc statistical analysis using the Tukey’s test showed that there were significant differences between the negative control and the normal control groups (P value < 0.0001). In addition, there were significant differences between the high dose test (P < 0.0001) as well as the low dose test (P =0.0379) and the negative control groups. There were significant differences in liver weights between the normal control and the low dose test groups (P = 0.031). However, there were no significant differences between the high dose test and the normal control groups (P = 0.99). A graphical representation of the results is shown in figure 6.

**Figure 6.**
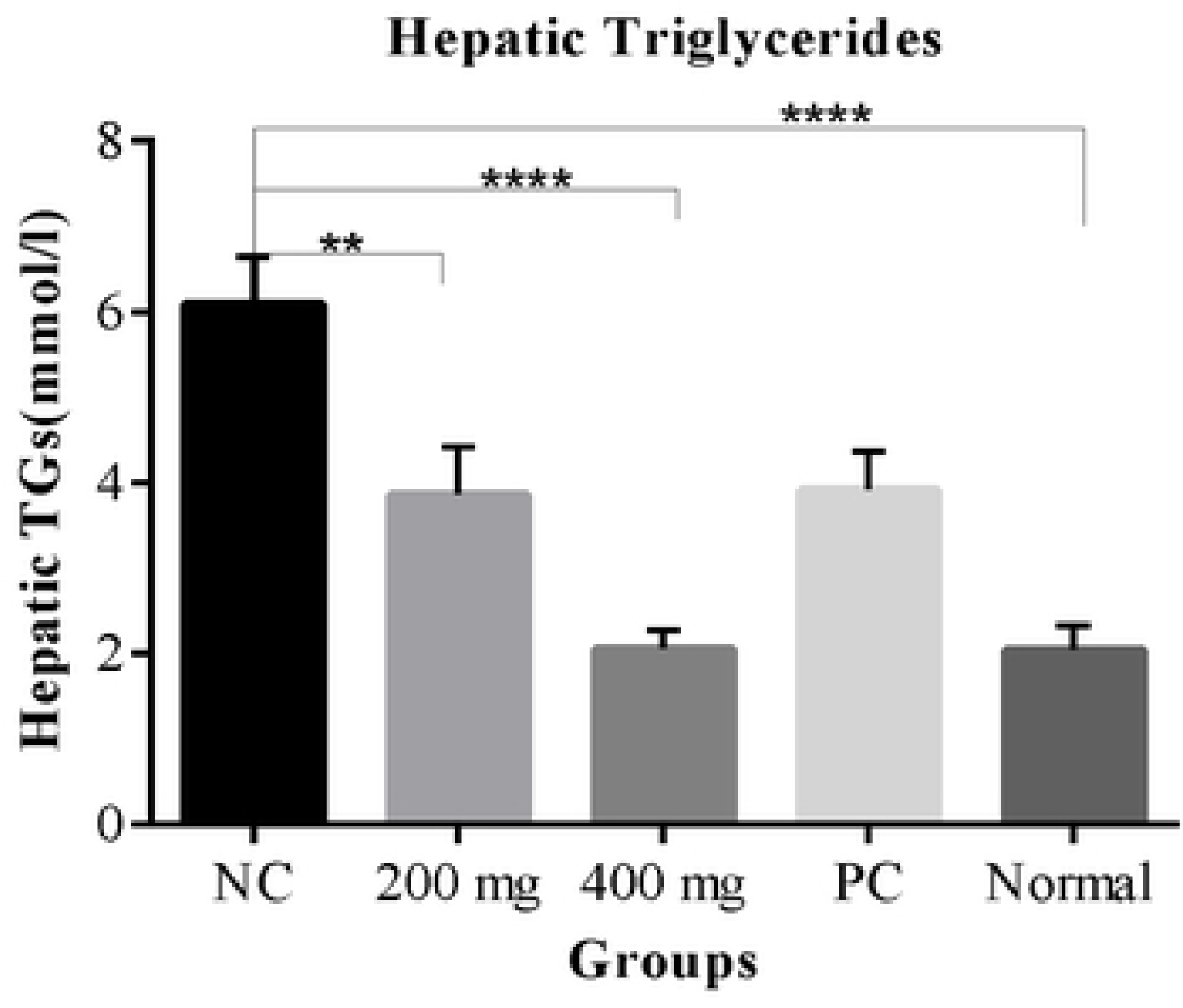
Mean liver weights in grams. NC (Negative control), PC (Positive control) * P value < 0.05, **P value < 0.01, **** P value < 0.0001.

### 3.6 Hepatic Index

There were significant differences in the hepatic index (mean liver-body weight ratio presented as a percentage (%)) between the experimental groups [6.1± 0.37% (negative control) vs. 2.4±0.17% (high dose test) vs. 3.88±0.37% (low dose test) vs. 4.4± 0.28% (positive control) vs. 1.7± 0.08% (normal control): P < 0.0001].

Post-hoc statistical analysis using the Tukey’s test showed that there were significant differences between the negative control and the high dose test (P < 0.0001) as well as the low dose test groups (P < 0.0001). There were significant differences between the high dose and low dose test groups (P value = 0.0022). There were significant differences between the low dose test and the normal control groups (P <0.0001). In contrast, there were no significant differences between the high dose test and the normal control groups (P value = 0.3874).

A graphical representation of the results is shown in figure 7.

**Figure 7.**
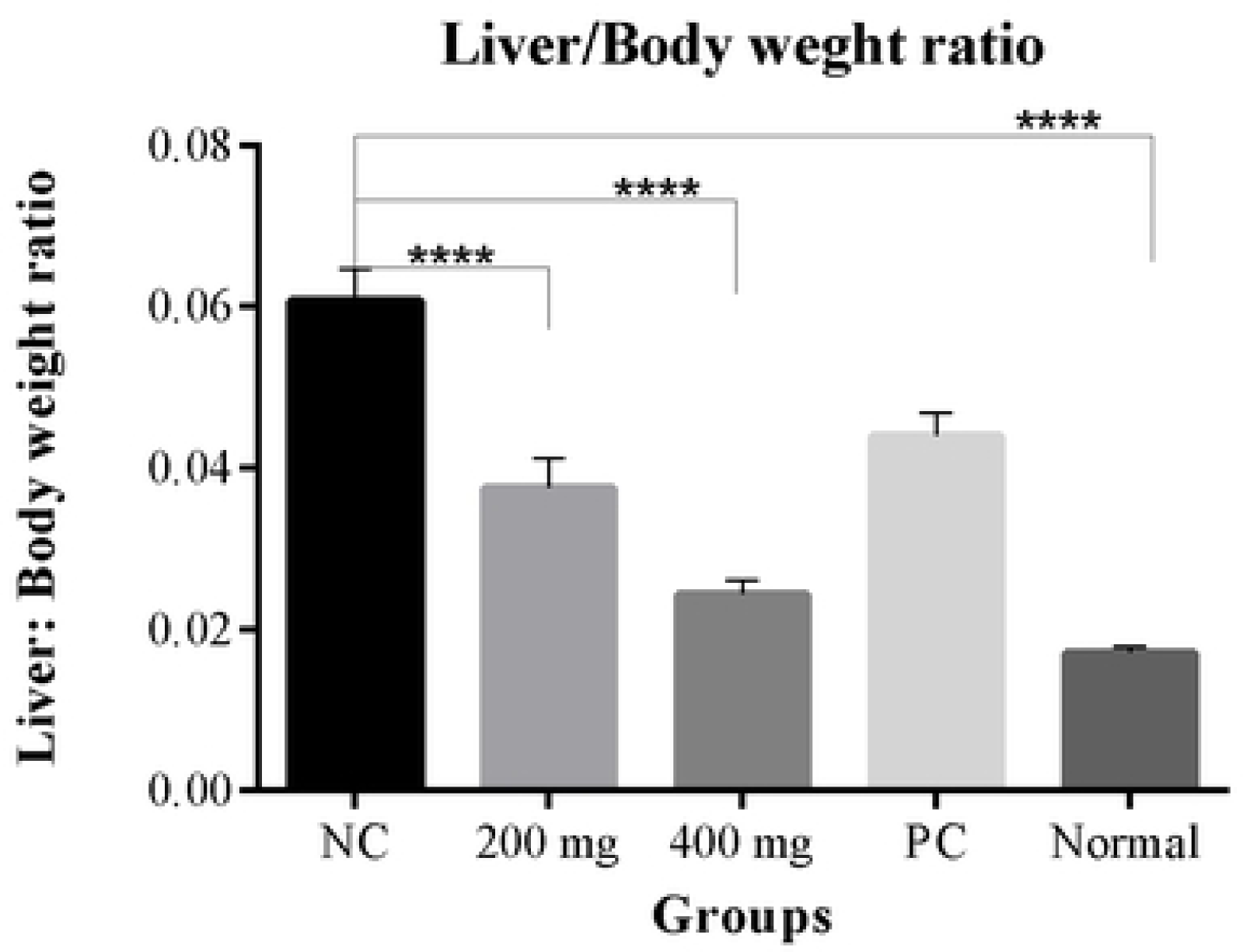
Hepatic index. NC (Negative control), PC (Positive control) ****P value < 0.0001.

### 3.7 Hepatic Function Tests

#### 3.7.1 Serum ALT

There were significant differences in serum ALT concentration between the five groups [155.6 ± 4.59 IU (negative control) vs. 91 ± 6.85 IU (high dose test) vs. 116.1 ± 3.96 IU (low dose test) vs. 119.9 ± 9.02 IU (Positive control) vs. 76.38 ± 5.01 IU (normal control): P value < 0.0001]. Post-hoc statistical analysis using the Tukey’s test showed there were significant differences between the negative control and the normal control groups (P < 0.0001). There were also significant differences between the high dose test and negative control groups (P < 0.0001) as well as between the Low dose test and negative control groups (P <0.006). There were also significant differences between the high dose and low dose test groups (P = 0.0493). A graphical representation of the results is shown in figure 8.

**Figure 8.**
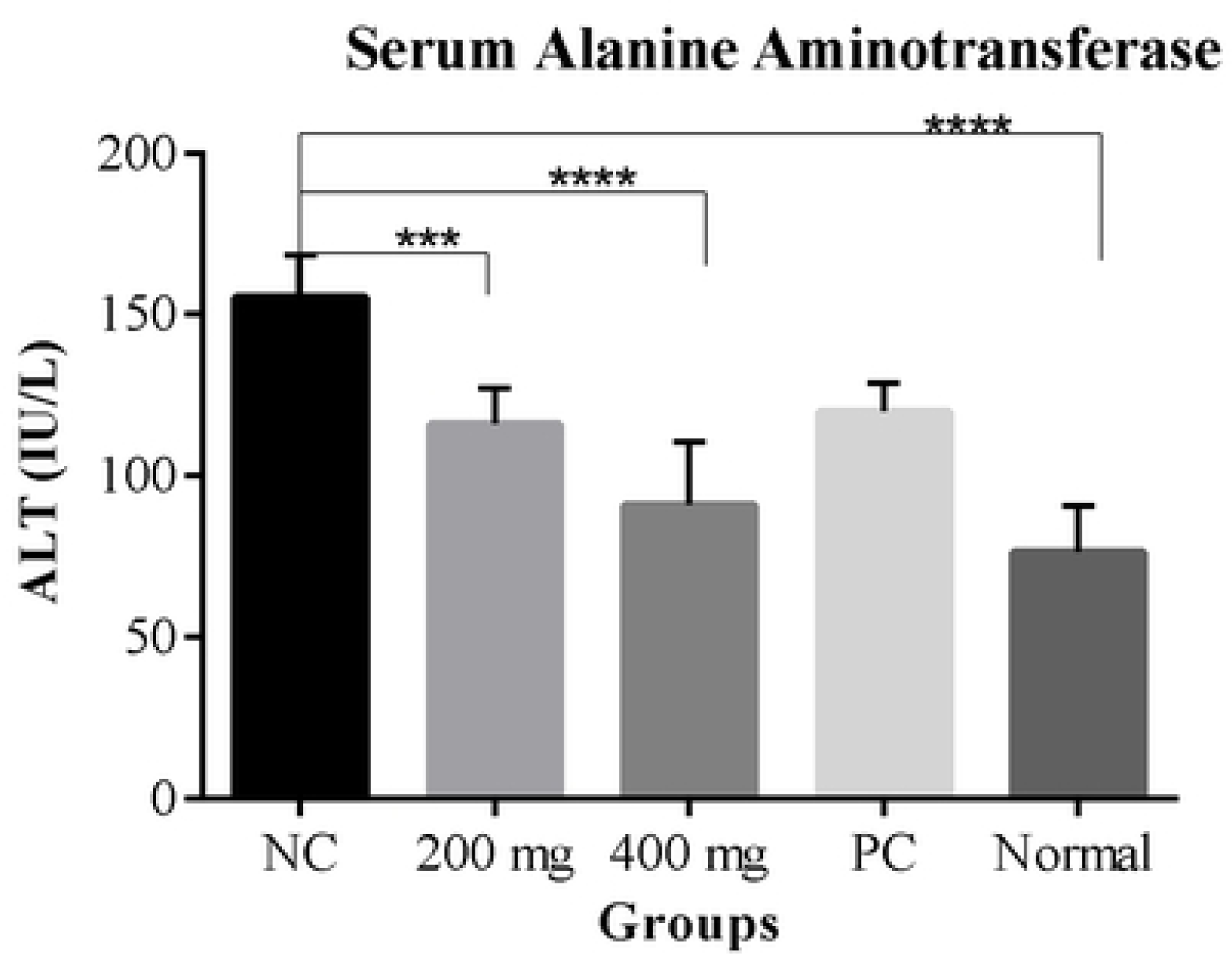
serum ALT in IU/L NC (Negative control), PC (Positive control). *P value < 0.05, *** P value < 0.001, **** P value < 0.0001.

#### 3.7.2 Serum AST levels

There were significant differences in serum AST levels between the five groups [133 ± 8.27 IU/L (low dose test) vs. 123.6 ± 11.72 IU/L (high dose test) vs. 150.9 ± 8.39 IU/L (negative control) vs. 132.8 ± 11.89 IU/L (positive control) vs. 92.7 ± 4.05 IU/L (normal control): P < 0.0017]. Post-hoc statistical analysis using the Tukey’s test showed significant differences between the normal control and low dose test groups (P = 0.0294). There were significant differences between the high dose test and the normal control groups (P = 0.308).

There were however no significant differences between the negative control and the low dose test groups (P = 0.648) as well between the negative control and the high dose test groups (P = 0.243). A graphical representation of these results is shown in figure 9.

**Figure 9.**
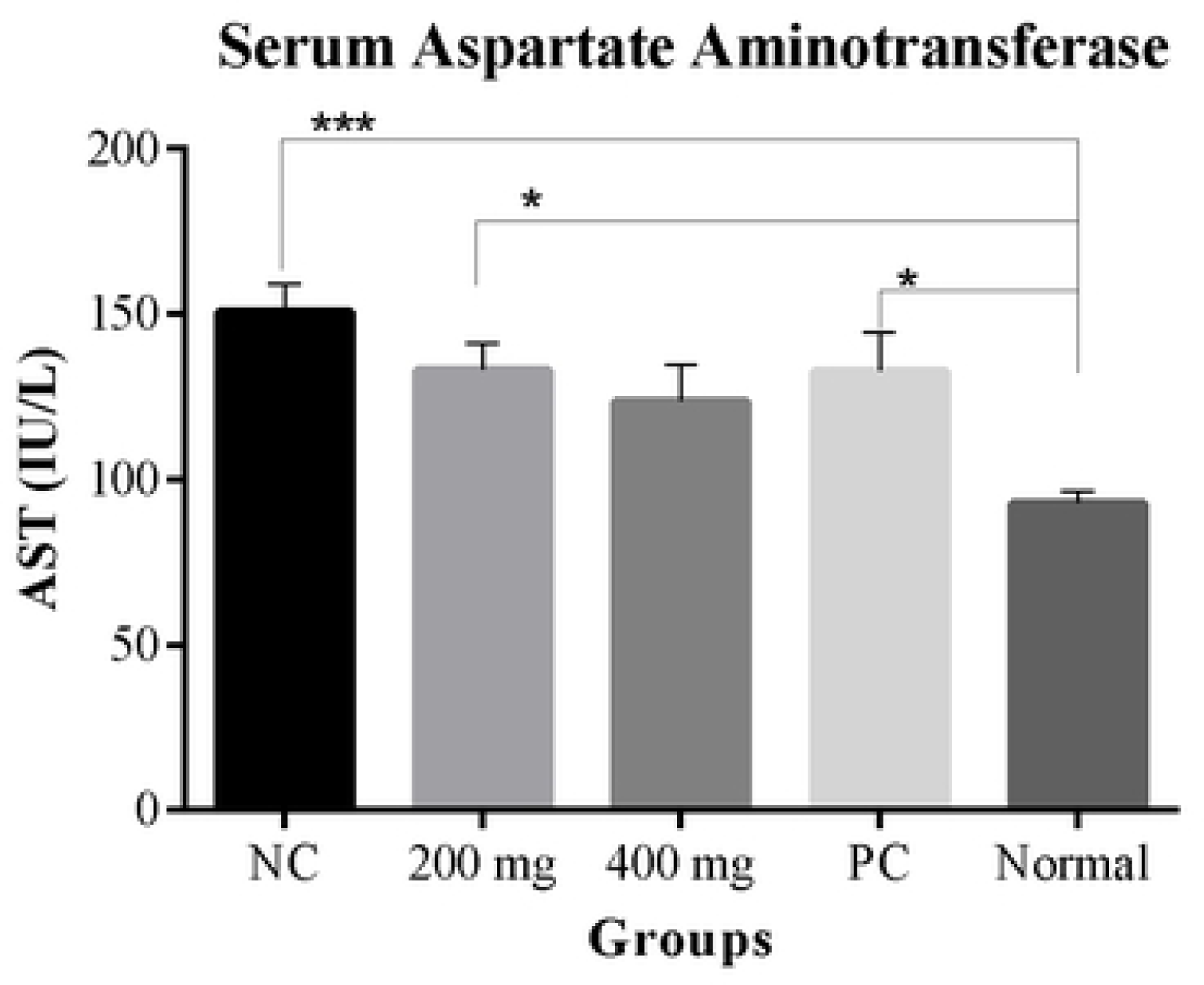
serum AST in IU/L. NC (Negative control), PC (Positive control). *p < 0.05, ***P < 0.00001.

### 3.8 Lipid Profiles

#### 3.8.1 Total Cholesterol

There were significant differences in serum total cholesterol levels between the five groups [3.39 ± 0.21 mmol/L (negative control) vs. 1.91 ± 0.16 mmol/L (high dose test) vs. 2.58 ± 0.15 mmol/L (low dose test) vs. 2.59 ± 0.16 mmol/L (positive control) vs. 1.62 ± 0.15 mmol/L (normal control): P < 0.0001].

Post-hoc statistical analysis using the Tukey’s test showed that there were significant differences between the negative control and the normal control groups (P < 0.0001). There were also significant differences between the negative control and the high dose test groups (P < 0.0001) as well as between the negative control and the low dose test groups (P = 0.0071). There were also significant differences between the high dose and low dose test groups (P = 0.0419).

A graphical representation of the results is shown in Fig. 10.

**Figure 10.**
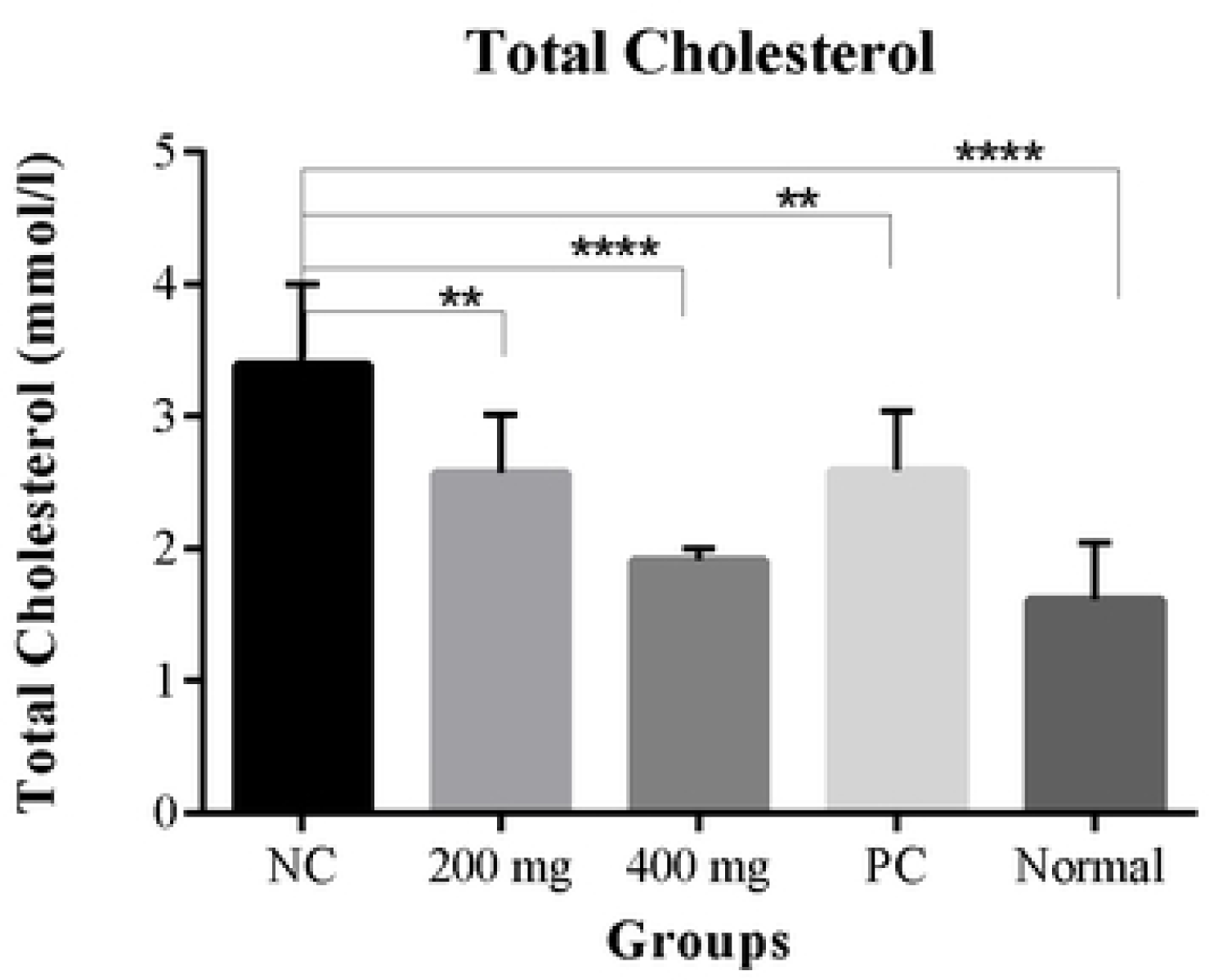
of serum total cholesterol in mmol/l. NC (Negative control), PC (Positive control) *P value < 0.05, **P value < 0.01, ***P value < 0.001, **** P value < 0.0001.

#### 3.8.2 Serum triglycerides

There were significant differences in the serum triglyceride levels among the five groups [2.20 ± 0.16 mmol/L (negative control) vs. 0.74 ± 0.12 mmol/L (high dose test) vs. 1.26 ± 0.16 mmol/L (low dose test) vs. 1.99 ± 0.07 mmol/L (positive control) vs. 0.5 ± 0.06 mmol/L (normal control): P < 0.0001].

Post-hoc statistical analysis using the Tukey’s test showed that there were significant differences between the negative control and the high dose test groups (P < 0.0001) as well as between the low dose test and negative control groups (P < 0.0001). There were also significant differences between the high dose and low dose test groups (P = 0.0345).

A graphical representation of these results is shown Figure 11.

**Figure 11.**
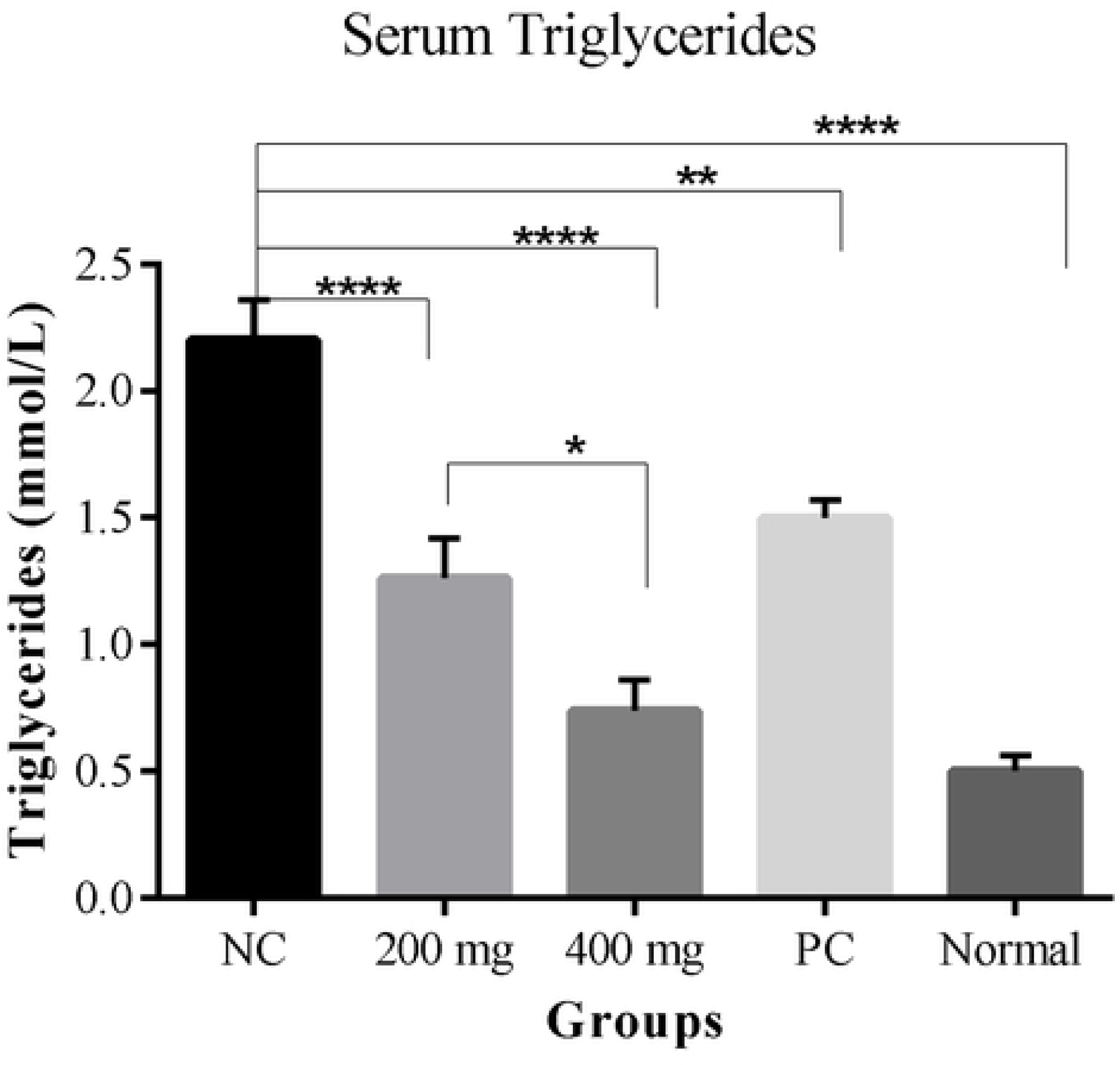
Serum triglyceride levels in mmol/L. NC (Negative control), PC (Positive control). *P value < 0.05, ****P value < 0.0001.

#### 3.8.3 Serum LDL cholesterol

There were significant differences in serum LDL cholesterol concentration among the five groups [2.02 ± 0.27 mmol/L (negative control) vs. 0.56 ± 0.09 mmol/L (high dose test) vs. 1.30 ± 0.11 (low dose test) vs. 1.16 ± 0.14 mmol/L (positive control) vs. 0.05 ± 0.08 mmol/L (normal control): P < 0.0001].

Post-hoc statistical analysis using the Tukey’s test showed that there were significant differences between the negative control and high dose test groups (P < 0.0001) as well as between the low dose test and negative control groups (P = 0.0172). In addition, there were significant differences between the low dose test and high dose test groups (P = 0.0149).

A graphical representation of these results is represented in Fig. 12.

**Figure 12.**
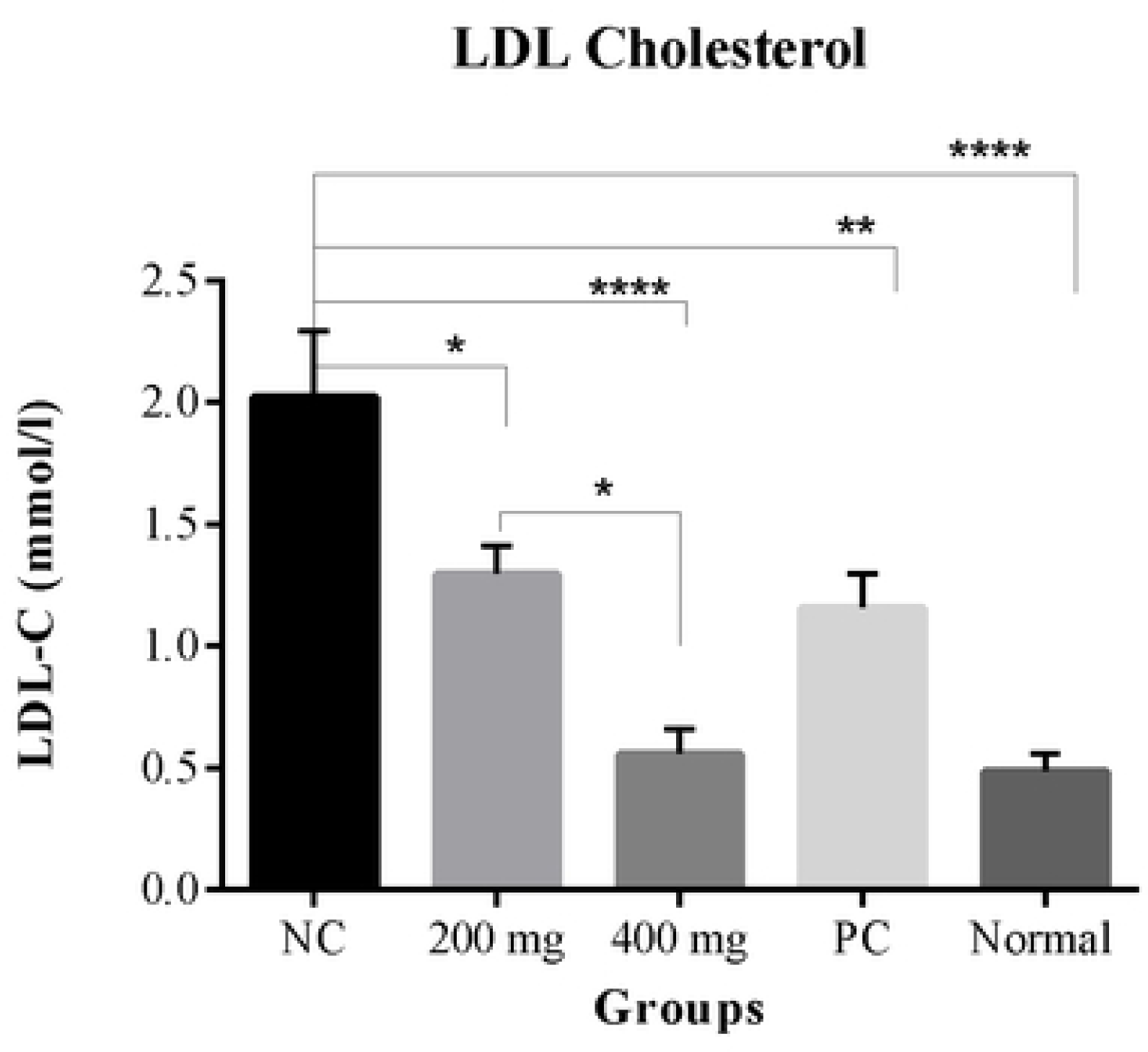
Serum LDL-cholesterol in mmol/L. NC (Negative control), PC (Positive control) * P value < 0.05, **P value <0.01, ****P value < 0.0001.

There were no statistically significant differences in HDL cholesterol concentration among the five groups (P value = 0.4759). However, the negative control group had the lowest HDL concentration among the groups

### 3.9 Histological evaluation of the liver

There were significant differences in hepatocellular ballooning among the five groups (P < 0.0001). Post-hoc analysis statistical demonstrated that the negative control group had significantly higher hepatocellular ballooning than the normal control group (P < 0.0001). There were significant differences between the negative control and the high dose test group (P = 0.0002 as well as between the low dose test and negative control groups (P = 0.0055). These results are represented in table 2 and table 3.

**Table 1.** Table showing incidence rates (%) of steatosis and lobular inflammation among the different groups.

|  | **STEATOSIS** |  |  |  | **LOBULAR INFLAMMATION** |  |  |  |
| --- | --- | --- | --- | --- | --- | --- | --- | --- |
| **GRADE** | 0 | 1 | 2 | 3 | 0 | 1 | 2 | 3 |
| **200mg/kg** | 62.5% | 25% | 12.5% | — | 75% | 25% | — | — |
| **400mg/kg** | 87.5% | 12.5% | — | — | 100% | — | — | — |
| **Negative control** | — | 25% | 75% | — | — | 62.5% | 37.5% | — |
| **Positive control** | 75% | 25% | — | — | 75% | 25% | — | — |
| **Normal control** | 100% | — | — | — | 100% | — | — | — |

**Table 2.**
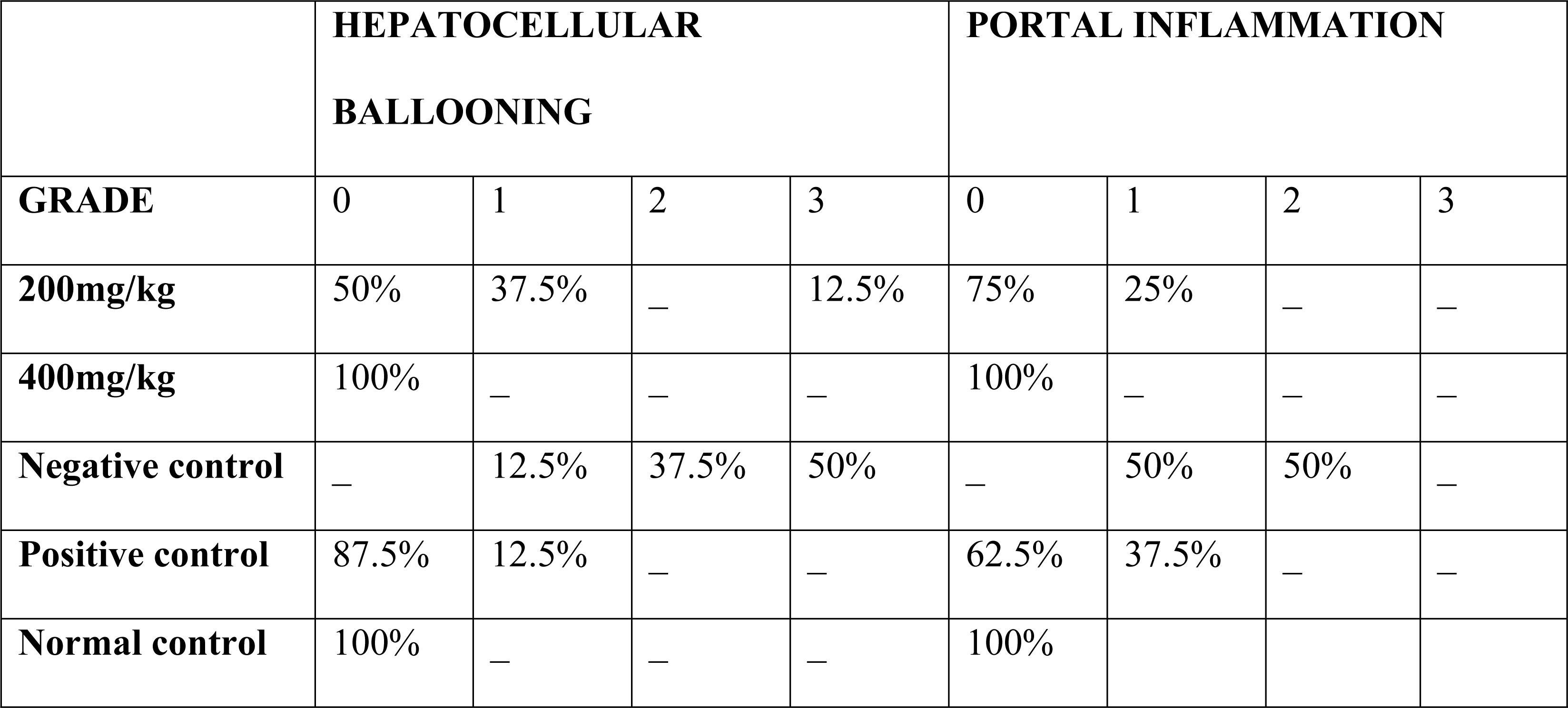
Table showing incidence rates (%) of hepatocellular ballooning and portal inflammation.

**Table 3.** Histology data presented as median (min.-max.) *significantly different from the Negative control group (p value< 0.05).

| **HISTOLOGICAL**<br>**FEATURE** | **200mg/kg** | **400mg/kg** | **Negative**<br>**control** | **Positive**<br>**control** | **Normal**<br>**control** |
| --- | --- | --- | --- | --- | --- |
| **Macro-vesicular**<br>**steatosis** | — | — | — | — | — |
| **Micro-vesicular**<br>**steatosis** | 0(0-1)* | 0(0-1)*** | 1.5(1-2) | 0(0-1)** | 0(0-0)*** |
| **Hepatocellular ballooning** | 0.5(0-1)** | 0(0-1)*** | 2.5(2-3) | 0.5(0-2)** | 0(0-0)**** |
| **Lobular inflammation** | 0(0-1)** | 0(0-1)*** | 0(1-2) | 0(0-1)** | 0(0-0)*** |
| **Portal inflammation** | 0(0-0) | 0(0-0) | 0.5(0-2) | 0(0-1) | 0(0-0) |

There were significant differences in occurrence of micro vesicular steatosis among the five groups (P < 0.0001). Post hoc statistical analysis showed that the negative control group had significantly more steatosis compared to: the high dose test (P = 0.0009), low dose test (P = 0.0009) and the normal control (P < 0.0001) groups. These results are represented in table 1 and table 3

There were significant differences in lobular inflammation among the 5 groups (P < 0.0001). Post-hoc statistical analysis showed that there were significant differences in lobular inflammation in the negative control group compared to: high dose test (P = 0.0001), low dose test (P = 0.0060) and normal control groups (P = 0.0001). These results are represented in table 1 and table 3.

There were no significant differences between the high dose test and the low dose test groups with respect to hepatocellular ballooning (P = 0.99), micro vesicular steatosis (P = 0.99) and lobular inflammation (P = 0.99).

There were significant differences in portal inflammation between the five groups (P value = 0.0460) but no significant differences were observed on post-hoc statistical analysis (P>0.05). The negative control group however had the highest incidence rates of grade 2 (50%) and grade 1 (50%) portal inflammation. These results are represented in table 2 and table 3.

There was no fibrosis or macro vesicular steatosis was observed in any of the experimental groups.

## 4.0 Discussion

Nonalcoholic fatty liver disease (NAFLD) covers a wide range of hepatic changes, ranging from simple steatosis to non-alcoholic steato-hepatitis (NASH), liver cirrhosis and hepatocellular carcinoma and is one of the most common chronic liver disorders worldwide [1,3,5]. It is associated with diverse clinical conditions including but not limited to obesity, renal failure, osteoporosis, type 2 diabetes, myocardial infarction, hypothyroidism, male infertility among others [5,6,8,21–25]. Current therapies for this condition however remain unsatisfactory underscoring the need for discovering new efficacious treatments for this condition [12,13,26].

Various animal models that aim to mirror both the histopathology and the pathophysiology of each stage of human NAFLD have been developed [27,28]. Indeed, selection of the appropriate animal model remains an important consideration in experimental design. We used a modified high fat/high sugar diet which was prepared by the addition of 0.8% monosodium-glutamate to a standard high-fat/high sugar diet. High-fat/high-sugar diets (“cafeteria diets”) are a popular animal model for the induction of NAFLD and are normally administered chronically over a period of six to sixteen weeks [29]. However, high-fat/high-sugar diets administered ad-libitum normally have the limitation of reduced food consumption compared to normal rat chow and this has been attributed to reduced palatability and satiety induction in the cafeteria diets [30]. Monosodium glutamate is a dietary tastant that confers the umami flavor to food and has been previously shown to increase food consumption was therefore added to the diet.

The NALFD model used in this study reproduced the serum biochemical and histological (steatosis, inflammation and hepatocellular injury) features of NAFLD [27] showing that the model possessed both construct and ecological validity and can therefore be used experimentally in the evaluation of anti-NAFLD therapies.

This study investigated the effects of the chronic administration of the freeze-dried stem bark extracts of *Erythrina abyssinica* on the development of NAFLD. This plant species finds extensive application in traditional medicine and is used in the preparation of health tonics by the Maasai community of East Africa [31,32]. In addition, this plant is shown to be rich in flavonoids, a group of natural products known to be efficacious in the management of NAFLD and other metabolic syndrome conditions [33–36].

The freeze-dried stem bark extracts of *Erythrina abyssinica* displayed potent dose-dependent antihyperglycemic effects in rats chronically fed a high-fat/high-sugar diet. Previous studies have shown that plant species belonging to this genus including the species that is the subject of this study possess antihyperglycemic activity [37,38]. The freeze-dried extracts of *Erythrina abyssinica* also possessed significant effects on insulin sensitivity. This was a significant finding in our opinion since insulin resistance is regarded as the most important pathophysiological mechanism of NAFLD [10,39]. In addition, it is also known that insulin resistance is one of the multiple hits determining the progression from NAFLD to non-alcoholic steato-hepatitis (NASH) [10,20,40]. It can therefore be seen that the extract can potentially prevent the establishment of, as well as slow the progression of NAFLD via the positive effects on insulin resistance.

There several putative mechanistic explanations for the observed effects on blood glucose levels as well as the insulin resistance. Benzo furans and Coumestan which possess agonist activities on AMP Kinase (AMPK) have been isolated from this plant species [41]. Activation of this enzyme is an attractive pharmacological target for the treatment of type 2 diabetes and other metabolic disorders [42]. Indeed leading diabetic drugs such as metformin and rosiglitazone mediate their metabolic effects partially through AMPK activation [42–44].

In addition, fifteen pterocarpan derivatives which possess inhibitory effects on protein tyrosine phosphatase 1B (PTP1B) have been isolated from the stem bark of this plant species [33,38]. Protein tyrosine phosphatase 1B (PTP1B) is a negative regulator of the leptin and insulin signaling pathways and plays important roles in the pathogenesis of obesity and diabetes [45]. The inhibition of this enzyme can therefore be expected to have beneficial effects in obesity, diabetes as well as NAFLD. This observation is supported by the fact that mice with the whole body deletion of PTP1B were protected against the development of obesity and diabetes [45,46].

The freeze-dried extracts possessed significant anti-dyslipidemic effects i.e. lowering plasma total-cholesterol, LDL-cholesterol and serum triglycerides. This was an important finding in our opinion. This is because NAFLD is characterized by atherogenic dyslipidemia, postprandial lipemia and high-density lipoprotein (HDL) dysfunction which are established risk factors for both cardiovascular and liver disease [47,48]. A possible mechanistic explanation for the observed anti-dyslipidemic affects is the AMP kinase inhibition described previously. It is known that AMP-kinase phosphorylates and inactivates the Sterol Regulatory Element Binding Protein (SREBP1) which has been implicated in the hepatic accumulation seen in NAFLD via its effects on lipid biosynthesis, LDL receptor biosynthesis and de novo cholesterol biosynthesis [49].

The freeze-dried stem bark extracts of *Erythrina abyssinica* significantly reduced hepatic lipid deposition as shown by the reduced hepatic triglyceride concentrations, liver weights and liver weight-body. These hepatic indices have been shown to be increased in NAFLD as well as in insulin resistance [50]. This was also a key finding in our opinion since according to the “two-hit” hypothesis on the pathophysiology of NAFLD, the hepatic triglyceride deposition is the first hit while the lipotoxicity that accompanies this deposition leads to hepatic injury and inflammation marks the second hit [51].

The extract of *Erythrina abyssinica* had significant effects on the serum alanine amino-transferase levels (ALT) which are a recognized marker of hepatocellular injury. Indeed, several epidemiologic association studies have shown that this enzyme is elevated in, and consequently can be used as a biomarker of NAFLD [52–54].

The extract significantly prevented the development of Histopathological evaluation of the liver showed prevention of steatosis, inflammation and hepatocellular injury. These effects were especially marked in the higher dose test group (400 mg/kg). It can therefore be seen that the results of the histopathological examination agree with those of the hepatic enzymes assay: i.e. that the extracts prevent the development of hepatic injury despite the chronic administration of a high fat/high sugar diet for eight (8) weeks.

### Conclusions and future directions

The freeze-dried extracts of *Erythrina abyssinica* had significant protective effects against the development of NAFLD in this study and show potential as a prophylactic against the development of this condition. These beneficial actions can be attributed to the rich variety of chemical compounds found in this plant species which have been shown to have positive effects in metabolic syndromes, cardiovascular disease and other conditions associated with aging [33,35,38,41,55]. The results of this study also validate the traditional use of this plant species as a tonic for the maintenance of good health.

## Acknowledgements

The authors would like to especially thank Patrick Mutiso Chalo from the University of Nairobi Herbarium for his assistance in the collection and identification of the plant species, Francis Okumu for the Department of Veterinary Anatomy and Physiology for his help in the microscopy section. The staff of the Department of Medical Physiology are also deeply thanked for their cooperation during this study.

## References

1. Sass DA, Chang P, Chopra KB. Nonalcoholic fatty liver disease: a clinical review. Dig Dis Sci. 2005;50: 171–80. Available: http://www.ncbi.nlm.nih.gov/pubmed/15712657

2. Masuoka HC, Chalasani N. Nonalcoholic fatty liver disease: an emerging threat to obese and diabetic individuals. Ann N Y Acad Sci. 2013;1281: 106–122. doi:10.1111/nyas.12016

3. Chalasani N, Younossi Z, Lavine JE, Charlton M, Cusi K, Rinella M, et al. The diagnosis and management of nonalcoholic fatty liver disease: Practice guidance from the American Association for the Study of Liver Diseases. Hepatology. 2018;67: 328–357. doi:10.1002/hep.29367

4. Chalasani N, Younossi Z, Lavine JE, Diehl AM, Brunt EM, Cusi K, et al. The diagnosis and management of non-alcoholic fatty liver disease: Practice Guideline by the American Association for the Study of Liver Diseases, American College of Gastroenterology, and the American Gastroenterological Association. Hepatology. 2012;55: 2005–2023. doi:10.1002/hep.25762

5. Byrne CD, Targher G. NAFLD: A multisystem disease. J Hepatol. 2015;62: S47–S64. doi:10.1016/j.jhep.2014.12.012

6. Mantovani A, Dauriz M, Gatti D, Viapiana O, Zoppini G, Lippi G, et al. Systematic review and meta-analysis: non-alcoholic fatty liver disease is associated with a history of osteoporotic fractures but not with low bone mineral density. Aliment Pharmacol Ther. John Wiley & Sons, Ltd (10.1111); 2019; doi:10.1111/apt.15087

7. Byrne CD, Perseghin G. Non-alcoholic fatty liver disease: A risk factor for myocardial dysfunction? J Hepatol. Elsevier; 2017;68: 640–642. doi:10.1016/j.jhep.2017.12.002

8. Mantovani A, Byrne CD, Bonora E, Targher G. Nonalcoholic Fatty Liver Disease and Risk of Incident Type 2 Diabetes: A Meta-analysis. Diabetes Care. 2018;41: 372–382. doi:10.2337/dc17-1902

9. Marchesini G, Mazzotti A. NAFLD incidence and remission: Only a matter of weight gain and weight loss? J Hepatol. 2015;62: 15–17. doi:10.1016/j.jhep.2014.10.023

10. Marchesini G, Bugianesi E, Forlani G, Cerrelli F, Lenzi M, Manini R, et al. Nonalcoholic fatty liver, steatohepatitis, and the metabolic syndrome. Hepatology. 2003;37: 917–923. doi:10.1053/jhep.2003.50161

11. Lonardo A, Targher G. From a fatty liver to a sugary blood. Dig Liver Dis. W.B. Saunders; 2018;50: 378–380. doi:10.1016/J.DLD.2018.01.126

12. Sridharan K, Sivaramakrishnan G, Sequeira RP, Elamin A. Pharmacological interventions for non-alcoholic fatty liver disease: a systematic review and network meta-analysis. Postgrad Med J. The Fellowship of Postgraduate Medicine; 2018;94: 556–565. doi:10.1136/POSTGRADMEDJ-2018-135967

13. Cholankeril R, Patel V, Perumpail B, Yoo E, Iqbal U, Sallam S, et al. Anti-Diabetic Medications for the Pharmacologic Management of NAFLD. Diseases. Multidisciplinary Digital Publishing Institute; 2018;6: 93. doi:10.3390/diseases6040093

14. Bellisle F. Effects of monosodium glutamate on human food palatability. Ann N Y Acad Sci. 1998;855: 438–41. Available: http://www.ncbi.nlm.nih.gov/pubmed/9929637

15. Brønstad A, Newcomer CE, Decelle T, Everitt JI, Guillen J, Laber K. Current concepts of Harm-Benefit Analysis of Animal Experiments - Report from the AALAS-FELASA Working Group on Harm-Benefit Analysis - Part j1. Lab Anim. SAGE Publications; 2016;50: 1–20. doi:10.1177/0023677216642398

16. Mandal J, Parija SC. Ethics of involving animals in research. Trop Parasitol. Wolters Kluwer -- Medknow Publications; 2013;3: 4–6. doi:10.4103/2229-5070.113884

17. Lee G, Goosens KA. Sampling blood from the lateral tail vein of the rat. J Vis Exp JoVE. 2015; e52766. doi:10.3791/52766

18. Bowe JE, Franklin Z, Hauge-Evan A, King A, Persaud S, Jones P. Metabolic phenotyping guidelines: assessing glucose homeostasis in rodent models. J Endocrinol. 2014;222: G13–G25. doi:10.1530/JOE-14-0182

19. Butler WM, Maling HM, Horning MG, Brodie BB. The direct determination of liver triglycerides. J Lipid Res. 1961;2: 95–96.

20. Kleiner DE, Makhlouf HR. Histology of Nonalcoholic Fatty Liver Disease and Nonalcoholic Steatohepatitis in Adults and Children. Clin Liver Dis. NIH Public Access; 2016;20: 293–312. doi:10.1016/j.cld.2015.10.011

21. Targher G, Byrne CD, Lonardo A, Zoppini G, Barbui C. Non-alcoholic fatty liver disease and risk of incident cardiovascular disease: A meta-analysis. J Hepatol. 2016;65: 589–600. doi:10.1016/j.jhep.2016.05.013

22. Pardhe BD, Shakya S, Bhetwal A, Mathias J, Khanal PR, Pandit R, et al. Metabolic syndrome and biochemical changes among non-alcoholic fatty liver disease patients attending a tertiary care hospital of Nepal. BMC Gastroenterol. BioMed Central; 2018;18: 109. doi:10.1186/s12876-018-0843-6

23. Guo Z, Li M, Han B, Qi X. Association of non-alcoholic fatty liver disease with thyroid function: A systematic review and meta-analysis. Dig Liver Dis. W.B. Saunders; 2018;50: 1153–1162. doi:10.1016/J.DLD.2018.08.012

24. Li Y, Liu L, Wang B, Chen D, Wang J. Nonalcoholic fatty liver disease and alteration in semen quality and reproductive hormones. Eur J Gastroenterol Hepatol. 2015;27: 1069–1073. doi:10.1097/MEG.0000000000000408

25. López-Lemus UA, Garza-Guajardo R, Barboza-Quintana O, Rodríguez-Hernandez A, García-Rivera A, Madrigal-Pérez VM, et al. Association Between Nonalcoholic Fatty Liver Disease and Severe Male Reproductive Organ Impairment (Germinal Epithelial Loss): Study on a Mouse Model and on Human Patients. Am J Mens Health. SAGE Publications; 2018;12: 639–648. doi:10.1177/1557988318763631

26. Bril F, Kalavalapalli S, Clark VC, Lomonaco R, Soldevila-Pico C, Liu I-C, et al. Response to Pioglitazone in Patients With Nonalcoholic Steatohepatitis With vs Without Type 2 Diabetes. Clin Gastroenterol Hepatol. 2018;16: 558–566.e2. doi:10.1016/j.cgh.2017.12.001

27. Lau JKC, Zhang X, Yu J. Animal models of non-alcoholic fatty liver disease: current perspectives and recent advances. J Pathol. Wiley-Blackwell; 2017;241: 36–44. doi:10.1002/path.4829

28. Takahashi Y, Soejima Y, Fukusato T. Animal models of nonalcoholic fatty liver disease/nonalcoholic steatohepatitis. World J Gastroenterol. Baishideng Publishing Group Inc; 2012;18: 2300–8. doi:10.3748/wjg.v18.i19.2300

29. Maeda Júnior A, Constantin J, Utsunomiya K, Gilglioni E, Gasparin F, Carreño F, et al. Cafeteria Diet Feeding in Young Rats Leads to Hepatic Steatosis and Increased Gluconeogenesis under Fatty Acids and Glucagon Influence. Nutrients. Multidisciplinary Digital Publishing Institute; 2018;10: 1571. doi:10.3390/nu10111571

30. Panchal SK, Brown L. Rodent models for metabolic syndrome research. J Biomed Biotechnol. Hindawi; 2011;2011: 351982. doi:10.1155/2011/351982

31. Parker ME, Chabot S, Ward BJ, Johns T. Traditional dietary additives of the Maasai are antiviral against the measles virus. J Ethnopharmacol. Elsevier; 2007;114: 146–152. doi:10.1016/J.JEP.2007.06.011

32. Kipkore W, Wanjohi B, Rono H, Kigen G. A study of the medicinal plants used by the Marakwet Community in Kenya. J Ethnobiol Ethnomed. BioMed Central; 2014;10: 24. doi:10.1186/1746-4269-10-24

33. Yenesew A, Twinomuhwezi H, Kiremire B, Mbugua M, Gitu P, Heydenreich M, et al. 8-Methoxyneorautenol and radical scavenging flavonoids from Erythrina abyssinica. Bull Chem Soc Ethiop. Chemical Society of Ethiopia; 2009;23. doi:10.4314/bcse.v23i2.44963

34. Yenesew A, Induli M, Derese S, Midiwo JO, Heydenreich M, Peter MG, et al. Anti-plasmodial flavonoids from the stem bark of Erythrina abyssinica. Phytochemistry. Pergamon; 2004;65: 3029–3032. doi:10.1016/J.PHYTOCHEM.2004.08.050

35. Prasain JK, Carlson SH, Wyss JM. Flavonoids and age-related disease: risk, benefits and critical windows. Maturitas. NIH Public Access; 2010;66: 163–71. doi:10.1016/j.maturitas.2010.01.010

36. Galleano M, Calabro V, Prince PD, Litterio MC, Piotrkowski B, Vazquez-Prieto MA, et al. Flavonoids and metabolic syndrome. Ann N Y Acad Sci. 2012;1259: 87–94. doi:10.1111/j.1749-6632.2012.06511.x

37. Kumar A, Lingadurai S, Shrivastava TP, Bhattacharya S, Haldar PK. Hypoglycemic activity of *Erythrina variegata* leaf in streptozotocin-induced diabetic rats. Pharm Biol. 2011;49: 577–582. doi:10.3109/13880209.2010.529615

38. Nguyen PH, Le TVT, Thuong PT, Dao TT, Ndinteh DT, Mbafor JT, et al. Cytotoxic and PTP1B inhibitory activities from Erythrina abyssinica. Bioorg Med Chem Lett. 2009;19: 6745–6749. doi:10.1016/j.bmcl.2009.09.108

39. Saponaro C, Gaggini M, Gastaldelli A. Nonalcoholic Fatty Liver Disease and Type 2 Diabetes: Common Pathophysiologic Mechanisms. Curr Diab Rep. 2015;15: 34. doi:10.1007/s11892-015-0607-4

40. Manco M. Insulin Resistance and NAFLD: A Dangerous Liaison beyond the Genetics. Child (Basel, Switzerland). Multidisciplinary Digital Publishing Institute (MDPI); 2017;4. doi:10.3390/children4080074

41. Nguyen P-H, Nguyen T-N-A, Dao T-T, Kang H-W, Ndinteh D-T, Mbafor J-T, et al. AMP-Activated Protein Kinase (AMPK) Activation by Benzofurans and Coumestans Isolated from *Erythrina abyssinica*. J Nat Prod. American Chemical Society and American Society of Pharmacognosy; 2010;73: 598–602. doi:10.1021/np900745g

42. Misra P, Chakrabarti R. The role of AMP kinase in diabetes. Indian J Med Res. 2007;125: 389–98. Available: http://www.ncbi.nlm.nih.gov/pubmed/17496363

43. Coughlan KA, Valentine RJ, Ruderman NB, Saha AK. AMPK activation: a therapeutic target for type 2 diabetes? Diabetes Metab Syndr Obes. Dove Press; 2014;7: 241–53. doi:10.2147/DMSO.S43731

44. Hardie DG. AMPK: a target for drugs and natural products with effects on both diabetes and cancer. Diabetes. American Diabetes Association; 2013;62: 2164–72. doi:10.2337/db13-0368

45. Panzhinskiy E, Ren J, Nair S. Pharmacological Inhibition of Protein Tyrosine Phosphatase 1B: A Promising Strategy for the Treatment of Obesity and Type 2 Diabetes Mellitus. Bentham Science Publishers; Available: https://www.ingentaconnect.com/content/ben/cmc/2013/00000020/00000021/art00001

46. Cho H. Protein Tyrosine Phosphatase 1B (PTP1B) and Obesity. Vitamins and hormones. 2013. pp. 405–424. doi:10.1016/B978-0-12-407766-9.00017-1

47. Peng K, Mo Z, Tian G. Serum Lipid Abnormalities and Nonalcoholic Fatty Liver Disease in Adult Males. Am J Med Sci. 2017;353: 236–241. doi:10.1016/j.amjms.2017.01.002

48. Chatrath H, Vuppalanchi R, Chalasani N. Dyslipidemia in Patients with Nonalcoholic Fatty Liver Disease. Semin Liver Dis. 2012;32: 022–029. doi:10.1055/s-0032-1306423

49. Kohjima M, Higuchi N, Kato M, Kotoh K, Yoshimoto T, Fujino T, et al. SREBP-1c, regulated by the insulin and AMPK signaling pathways, plays a role in nonalcoholic fatty liver disease. Int J Mol Med. 2008;21: 507–11. Available: http://www.ncbi.nlm.nih.gov/pubmed/18360697

50. Arab JP, Arrese M, Trauner M. Recent Insights into the Pathogenesis of Nonalcoholic Fatty Liver Disease. Annu Rev Pathol Mech Dis. Annual Reviews; 2018;13: 321–350. doi:10.1146/annurev-pathol-020117-043617

51. Buzzetti E, Pinzani M, Tsochatzis EA. The multiple-hit pathogenesis of non-alcoholic fatty liver disease (NAFLD). Metabolism. 2016;65: 1038–1048. doi:10.1016/j.metabol.2015.12.012

52. Khosravi S, Alavian SM, Zare A, Daryani NE, Fereshtehnejad S-M, Daryani NE, et al. Non-alcoholic fatty liver disease and correlation of serum alanin aminotransferase level with histopathologic findings. Hepat Mon. Kowsar Medical Institute; 2011;11: 452–8. Available: http://www.ncbi.nlm.nih.gov/pubmed/22087177

53. Sheng X, Che H, Ji Q, Yang F, Lv J, Wang Y, et al. The Relationship Between Liver Enzymes and Insulin Resistance in Type 2 Diabetes Patients with Nonalcoholic Fatty Liver Disease. Horm Metab Res. © Georg Thieme Verlag KG; 2018;50: 397–402. doi:10.1055/a-0603-7899

54. Schindhelm RK, Diamant M, Dekker JM, Tushuizen ME, Teerlink T, Heine RJ. Alanine aminotransferase as a marker of non-alcoholic fatty liver disease in relation to type 2 diabetes mellitus and cardiovascular disease. Diabetes Metab Res Rev. 2006;22: 437–443. doi:10.1002/dmrr.666

55. Majinda RRT, Wanjala CCW, Juma BF. Bioactive non-alkaloidal constituents from the genus Erythrina. Stud Nat Prod Chem. Elsevier; 2005;32: 821–853. doi:10.1016/S1572-5995(05)80070-5

